# Wavelet-Domain Multi-Representation and Ensemble Learning for Automated ECG Analysis

**DOI:** 10.64898/2026.02.14.705908

**Authors:** Lina Chato, Alex Kagozi

## Abstract

Accurate diagnosis of cardiac abnormalities from electrocardiogram signals remains a central challenge in automated cardiovascular assessment. This study investigates the efficiency of time–frequency representations and deep learning architectures in classifying 12-lead ECGs into five diagnostic super-classes using the PTB-XL dataset. Continuous Wavelet Transform is applied to generate time– frequency representations, scalograms and phasograms, representing spectral energy and phase distributions, respectively. We experiment with both early and late information fusion strategies using several convolutional and transformer-based networks of a custom Convolutional Neural Network, Hybrid Deep Learning, transfer learning, feature fusion, and ensemble modeling, and weighted loss strategies. An ensemble fusion of models trained on time-frequency representation and time representation achieved the best overall performance of Area Under Curve of 0.9233 surpassing individual modalities. To improve the results further, weighted focal loss is used to improve the low classification rates in some labels due to imbalanced data. The results highlight the potential of multi-representation wavelet fusion for interpretable and generalizable ECG classification.

## 1. INTRODUCTION

Cardiovascular diseases (CVDs) are the leading cause of death globally [1]. Electrocardiography (ECG) is the most common non-invasive diagnostic tool, yet manual interpretation is time-consuming and prone to variability [2], motivating automated ECG classification using deep learning (DL). Recent DL models, particularly convolutional and recurrent neural networks, have achieved high accuracy in arrhythmia and myocardial infarction detection [3–6]; however, most rely solely on time-domain waveforms and overlook the joint time–frequency information that characterizes pathological ECG patterns. The Continuous Wavelet Transform (CWT) provides a richer representation by capturing both temporal and spectral dynamics. Its magnitude (scalogram) and phase (phasogram) images allow DL models to interpret ECGs as structured inputs, improving robustness and leveraging pretrained image-based models. *Zyout et al*. applied scalograms of individual ECG waves with Convolutional Neural networs (CNNs) to distinguish normal, tachycardia, and bradycardia rhythms, achieving over 98% accuracy [7]. *Scarpiniti et al*. explored early, intermediate, and late fusion of scalogram and phasogram inputs to enhance arrhythmia classification [8].

The PTB-XL dataset is one of the largest publicly available 12-lead ECG datasets, covering a wide range of clinically relevant cardiac conditions with multi-class and multi-label annotations [9]. Its scale, diversity, and high-quality annotations make it an ideal benchmark for developing and evaluating DL models for automated ECG analysis. Despite this, to our knowledge, no study has yet explored the representations of scalogram, phasogram, or their combinations derived from the PTB-XL dataset for deep learning-based multi-class, multi-label classification, highlighting a gap that our work addresses.

In this research, we propose a wavelet-based multi-representation framework using scalograms, phasograms, and their combinations trained under early and late fusion strategies across several architectures, custom CNN, and transfer learning strategies. In the early fusion approach, scalograms and phasograms were concatenated channel-wise to form unified multimodal inputs, allowing the network to learn joint spectral–phase representations within a single feature space. In contrast, the late fusion strategy trained independent models for each modality, and their output logits combined through weighted averaging to generate the final prediction. Pretrained image-based models, hybrid models, and ensemble techniques have been proposed and examined to develop a robust multi-class multi-label ECG classification system.

The reminder of this paper is presented as follows: *Section 2* describes the data preparations, and the proposed models; *Section 3* discusses experiments and displays the results; Finally, *Section 4* concludes this paper and presents future research direction.

## 2. METHODOLOGY

### 2.1 Data Preprocessing

The PTB-XL dataset, curated by the Physikalisch-Technische Bundesanstalt (PTB) in collaboration with the Fraunhofer Heinrich Hertz Institute and Charité–Universitätsmedizin Berlin [9], contains 21,837 12-lead ECG recordings of 10 sec each, sampled at 500 Hz and 100 Hz. It defines several diagnostic tasks ranging from binary, normal vs. abnormal, to multi-label classification, including the Super-Diagnostic Task, which categorizes ECGs into five clinically relevant superclasses: Normal, Myocardial Infarction, ST/T Change, Conduction Disturbance, and Hypertrophy [10]. The data distribution of the main five classes is shown in **Figure 1**. However, each sample in the data may contain any combinations of the classes (i.e., labels) excluding the normal class. Thus, this research focuses on multi-class multilableing, leveraging multi-modal CWT representations to improve diagnostic accuracy and generalization. Following PTB-XL’s recommended stratified folds, data were partitioned into train, validation, and test with the ratio 8:1:1; the first eight folds for training (including 17,111 samples), the ninth fold for validation (including 2, 156 samples), and the tenth fold for test (including 2,163).

**Fig. 1.**
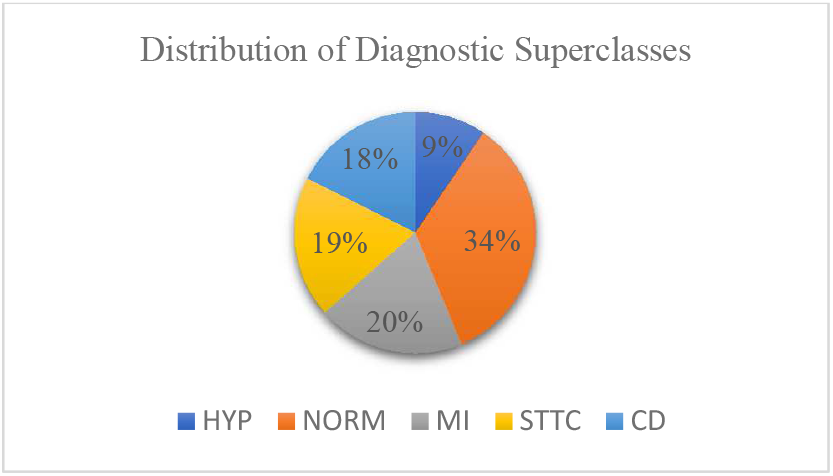
The distribution of PTB-XL dataset in terms of diagnostic super-classes. Where NORM: normal ECG, MI: myocardial infarction, CD: conduction disturbance, STTC: ST/T-changes, HYP: hypertrophy.

Three ECG signal processing methods are applied to prepare different representations of the data:

#### Band-pass filtering

Each ECG lead *X*_*i*_(*t*), where *i* = 1,2, …,12corresponds to one of the twelve standard ECG leads, was detrended and band-pass filtered within 0.05-47 Hz to suppress baseline wander and high-frequency noise as follows:

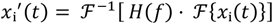

where ℱand ℱ^−1^denote the Fourier and inverse Fourier transforms, respectively, and *H*(*f*)represents the band-pass filter response.

Following filtering, **Z-score standardization** was applied to each lead to normalize the signals to zero mean and unit variance, given by

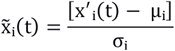

where *μ*_*i*_ and *σ*_*i*_ denote the mean and standard deviation of lead *i*, respectively.

#### Continuous Wavelet Transform

To extract time-frequency representations from each ECG signal (i.e., x(t)), the CWT is performed as follows:

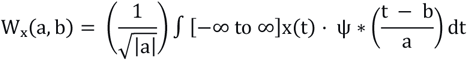

Where *a* represents the scale parameter (controls frequency resolution), *b* performs translation parameter (controls temporal shift), *ψ(t)* represents the Morlet wavelet, and * denotes complex conjugation.

From the complex *CWT* coefficients of *Wx(a,b)*, two distinct time–frequency representations are derived: the amplitude map (scalogram) and the phase map (phasogram), as shown in **Figure 2**. The scalogram captures localized power spectra using the squared magnitude of *Wx(a,b)*, followed by log-scaling and percentile-based clipping (1st–99th) to suppress outliers and enhance dynamic range. Both scalograms and phasograms were normalized per lead to the [0,1] range for numerical stability and resized to 224×224 pixels using bilinear interpolation. Each 12-lead ECG recording thus produced two corresponding image tensors, one for spectral amplitude and one for phase which served as model inputs.

**Fig. 2.**
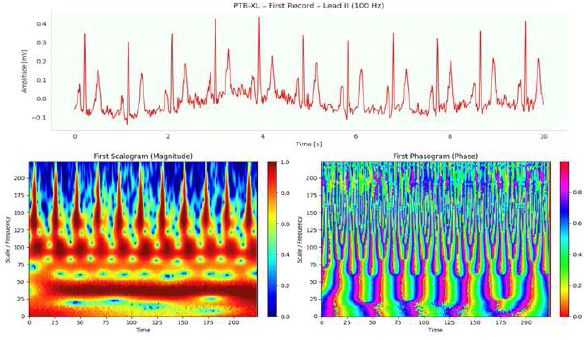
Sample ECG signal (lead II) (top) with corresponding CWT-based scalogram (left) and phasogram (right).

### 2.2. Multi-Class Multi-Label System Development

Our system integrates multiple modeling strategies across different data domains: time-domain raw ECG signals, timefrequency representations using CWT, and fusion of magnitude (scalogram) and phase (phasogram) features (early and late fusions). Additionally, pretrained imaging models based on transfer learning, and hybrid CNN– Transformer architectures are employed to enhance features quality, accelerate convergence, and improve multi-class multi-label classification performance.

#### 2.2.1 Time Domain – Raw Signals

As a baseline, we adopt the **XResNet1d101** from [3], a 1D variant of ResNet tailored for sequential ECG data. It includes a stem convolution (kernel size 7), four residual stages [3,4,23,3], and Rectifier Linear Unit (ReLU) activations with batch normalization. The network concludes with adaptive average pooling and a fully connected layer for 5-class multi-label classification.

#### 2.2.2 Time-Frequency Domain – Scalograms

Scalograms are processed as 12-channel 2D images (one per ECG lead). A custom ResNet-based CNN called CWT2DCNN. It is designed to handle 12-channel input, as shown in **Figure 3**. Additionally, pretrained models, such as Swin Transformer, ResNet50, and EfficientNetB0, are finetuned to improve performance and speed convergence. Since these models accept 3-channel 2D input, a 1×1 convolutional layer was applied to map the 12-lead scalograms to 3 channels, as shown in **Figure 4**. We also fine-tuned a hybrid CNN–Swin Transformer model in [10], using same strategy in **Figure 4**. All backbones (CWT2DCNN and pretrained models) were followed by an MLP of two Dense layers (512 and 256 units, respectively) and dropout 0.3, ending with a five-node outputs for multi-class multi-label classification.

**Fig. 3.**
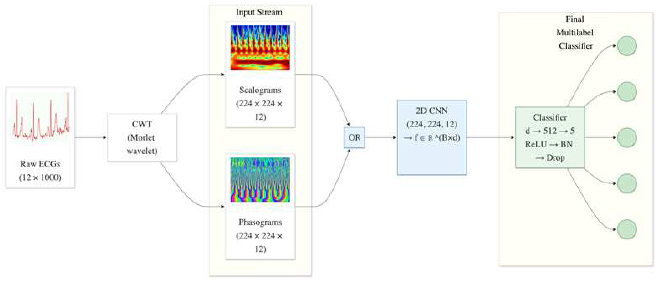
Input processing for scalogram only / or phasogram only features based on CWT2DCNN that uses all 12 channels natively from the specific CWT modality.

**Fig. 4.**
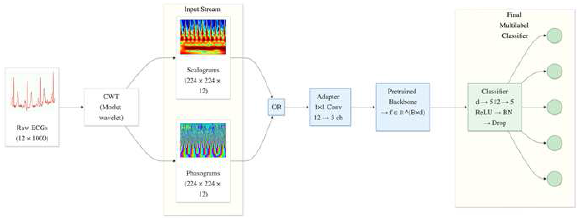
Input processing for scalogram only / or phasogram only features based on pretrained imaging models. The 12 channels from the specific used CWT modality are reduced to 3 channels to be compatible with the input of pretrained models.

#### 2.2.3 Time-Frequency Domain – Phasograms

The same architectures and strategies were described in the previous *subsection (2*.*2*.*2)*, are used to train on phasogram features (image of 12 channels, which are derived from the CWT phase components) instead of scalograms.

#### 2.2.4 Feature Fusion: Phasograms & Scalograms

To exploit complementary magnitude-phase information, we design **early** and **late fusion** strategies.

In **early fusion**, scalogram and phasogram tensors are concatenated into a 24-channel input, allowing joint spectral– phase learning. The CWT2DCNN is adapted to process 24 channels, as shown in **Figure 5**, while pretrained backbones are adapted via a 1×1 convolutional layer compressing 24 to 3 channels, as shown in **Figure 6**.

**Fig. 5.**
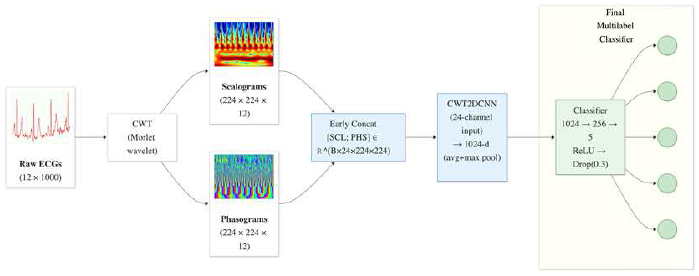
Early fusion of scalograms and phasograms. Inputs are concatenated into 24 channels. The Fusion-2DCNN processes all 24 channels.

**Fig. 6.**
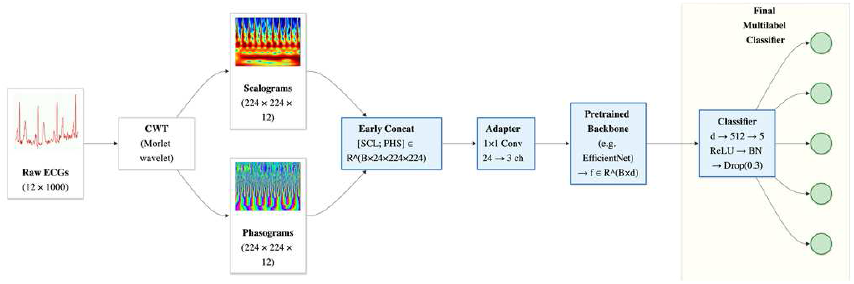
Early fusion of scalograms and phasograms. Inputs are concatenated into 24 channels.

In **late fusion**, each modality is processed through separate encoders, and their 512-dimensional embeddings are concatenated into a unified 1024-dimensional vector classified via an MLP, as shown in **Figure 7**. For pretrained models, dual encoders extract global features, fused through an MLP with dropout (p=0.3), as illustrated in **Figure 8**.

**Fig. 7.**
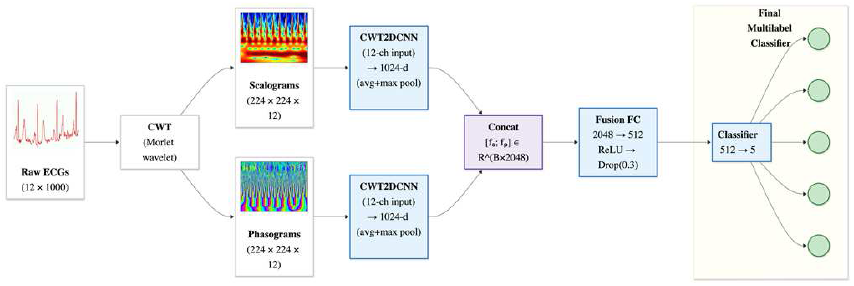
Late fusion architecture. Scalograms and phasograms.

**Fig. 8.**
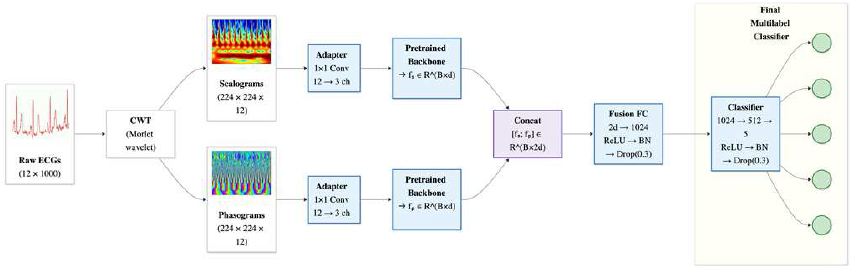
Late fusion architecture. Scalograms and phasograms are processed in parallel streams. Each stream supports pretrained 3-channel backbones (via adapter).

## 3. EXPERIMENTS AND RESULTS

All models were implemented in PyTorch 2.0 and trained with multi-label BCE or weighted Focal Loss to address class imbalance. Experiments were conducted on an Nvidia A100 GPU (40 GB) supported by Nautilus [11]. Performance was evaluated using macro AUC, F1, and Fβ (β = 2) on the PTB-XL patient-wise test split. Transformer-based models used batch size 16 and learning rate 5×10^−5^, while CNNs used batch size 32 with learning rates of 1×10^−3^–1×10^−4^. Training was performed for 50 epochs with early stopping (patience = 5). Table I reports BCE results and the best Focal Loss performance. The xresnet1d101 baseline achieved the best raw-signal performance (AUC = 0.9224). For wavelet-based representations, the hybrid Swin transformer performed best on both scalograms and phasograms, highlighting the benefit of hybrid CNN–Transformer feature extraction. Early fusion slightly outperformed late fusion, suggesting improved joint phase–amplitude learning. The ensemble of xresnet1d101 with early fusion hybrid Swin and ResNet50 achieved the top overall performance (AUC = 0.9238) using optimized focal loss weights. The code is available at: https://github.com/kagozi/MultiModal-EC

**TABLE I.** CLASSIFICATION PERFORMANCE ON THE PTB-XL DATASET IN TERMS OF MACRO AUC AND F1-Β FOR THE SUPER-DIAGNOSTIC TASK. GRAY-SHADED CELLS DENOTE MODELS TRAINED WITH WEIGHTED FOCAL LOSS, WHEREAS UNSHADED CELLS CORRESPOND TO **BCE** TRAINING. T_tr_ IS TRAIN TIME, AND T_ts_ TEST TIME.

| Domain | Model | AUC | $T_{ts}$<br>(ms) | $T_{tr}$<br>(hour) |
| --- | --- | --- | --- | --- |
| **Cross Entropy Loss** |  |  |  |  |
| Time (Raw Signals) | XResNet1d101 | **0.9224** | **1.412** | **0.42** |
| Time-Frequency Energy (Scalogram) | Swin Transformer | 0.8753 | 8.103 | 6.66 |
|  | ResNet50 | 0.8724 | 5.342 | 1.19 |
|  | Custom 2DCNN | 0.8724 | 4.880 | 0.46 |
|  | EfficientNet-B0 | 0.8693 | 5.173 | 1.34 |
|  | Hybrid Swin Transformer | 0.8819 | 7.578 | 4.17 |
| Time-Frequency Phase (Phasogram) | Swin Transformer | 0.8693 | 8.133 | 6.69 |
|  | ResNet50 | 0.8662 | 5.407 | 1.21 |
|  | Custom 2DCNN | 0.8475 | 4.908 | 0.48 |
|  | EfficientNet-B0 | 0.8516 | 5.183 | 1.36 |
|  | Hybrid Swin Transformer | 0.8712 | 7.536 | 4.19 |
| Early Fusion of Phasogram Scalogram | Swin Transformer | 0.8857 | 11.536 | 6.72 |
|  | ResNet50 | **0.8902** | **9.499** | **1.22** |
|  | Custom 2DCNN | 0.8818 | 8.287 | 0.54 |
|  | EfficientNet-B0 | 0.8727 | 10.173 | 1.47 |
|  | Hybrid Swin Transformer | **0.9018** | **11.503** | **4.41** |
| Late Fusion Phasogram Scalogram | Swin Transformer | 0.8807 | 15.928 | 13.31 |
|  | ResNet50 | 0.8832 | 9.261 | 2.39 |
|  | Dual Stream 2DCNN | 0.8810 | 8.287 | 0.97 |
|  | EfficientNet-B0 | 0.8719 | 10.834 | 2.71 |
|  | Hybrid Swin Transformer | 0.8839 | 13.422 | 8.28 |
| Ensemble (Best Three) |  | **0.9233** | **21.879** | - |
| **Weight Focal Loss weights = {1.0, 5.0, 2.5, 1.0, 0.5}** |  |  |  |  |
| Early Fusion | Hybrid Swin | **0.9023** | 11.536 | 4.35 |
| Early Fusion | ResNet50 | **0.8915** | 9.387 | 1.29 |
| Ensemble (best model) | The above 2 models+ XResNet1d101 | **0.9238** | 22.335 | - |

## 4. CONCLUSION AND FUTURE DIRECTIONS

This study proposed a DL framework for multi-class, multilabel ECG classification on the PTB-XL dataset. As summarized in Table I, wavelet-based feature fusion significantly improved performance over single-representation wavelet models, with both early and late fusion consistently providing gains. Early fusion achieved the better results than late fusion, indicating that joint phase– spectral learning enhances discriminative feature extraction. Weighted focal loss further boosted performance, by addressing class imbalance and improving minority-label detection. The highest overall accuracy was obtained through ensemble modeling combining the raw signal model (baseline) with the best early fusion hybrid architectures (Swin Transformer and ResNet50) outperformed all individual models and the baseline raw model [3].

In addition to accuracy, the framework demonstrated clinical feasibility in terms of efficiency. The raw model achieved 1.412 ms/sample inference time, while the hybrid Swin and ensemble models required only 11.536 ms/sample and 22.335 ms/sample, supporting potential real-time deployment. Although lightweight raw signal model (baseline) performed competitively, more complex structures based on feature fusion of wavelet components and ensemble modeling are better suited to capture subtle pathological patterns as larger and more balanced ECG datasets become available. Notably, *Dhyani et al*. employed the smaller MIT–BIH dataset with handcrafted wavelet features, but its beat-level two-lead setup and label mismatch make it unsuitable for augmentation in our PTB-XL multi-label diagnostic setting [12].

Future work will explore multi-head classification, generative ECG augmentation to further mitigate imbalance, and lead subset analysis for more efficient model designs.

## Notes

### Competing Interest Statement

The authors have declared no competing interest.

